# CancerVar: an Artificial Intelligence empowered platform for clinical interpretation of somatic mutations in cancer

**DOI:** 10.1101/2020.10.06.323162

**Authors:** Quan Li, Zilin Ren, Kajia Cao, Marilyn M. Li, Kai Wang, Yunyun Zhou

**Author notes:** To whom correspondence should be addressed: Dr. Kai Wang, Dr. Yunyun Zhou.

## Abstract

Several knowledgebases, such as CIViC and OncoKB, have been manually curated to support clinical interpretations of a limited number of “hotspot” somatic mutations in cancer, yet discrepancies or even conflicting interpretations have been observed among these knowledgebases. Additionally, while these knowledgebases have been extremely useful, they typically cannot interpret novel mutations, which may also have functional and clinical impacts in cancer. To address these challenges, we developed an automated interpretation tool called CancerVar (Cancer Variants interpretation) to score more than 12.9 million somatic mutations and classify them into four tiers: strong clinical significance, potential clinical significance, uncertain clinical significance, and benign/likely benign, based on the AMP/ASCO/CAP 2017 guideline. Considering that the AMP/ASCO/CAP rule-based scoring system may have inherent limitations, such as lack of a clear guidance on weighing different pieces of functional evidence or unclear definition for certain clinical evidence, it may cause misinterpretation for certain variants that have functional impacts but no proven clinical significance. To address this issue, we further introduced a deep learning-based scoring system to predict oncogenicity of mutations by semi-supervised generative adversarial network (SGAN) method using both functional and clinical evidence. We trained and validated the SGAN model on 5,234 somatic mutations from an in-house database of clinical reports on cancer patients, and achieved a good performance when testing on 6,226 variants that were curated by us through literature search. We also compared the prediction with several independent datasets and showed great utility in classifying variants with previously unknown interpretations. CancerVar is also incorporated into a web server that can generate automated texts with summarized descriptive interpretations, such as diagnostic, prognostic, targeted drug responses and clinical trial information for many hotspot mutations. In summary, CancerVar can facilitate clinical interpretation and hypothesis generation for somatic mutations, and greatly reduce manual workload for retrieving relevant evidence and implementing existing guidelines.

## INTRODUCTION

A large number of somatic variants have been identified by next-generation sequencing (NGS) during the practice of clinical oncology to facilitate precision medicine (1,2). In order to better understand the clinical impacts of somatic variants in cancer, several knowledgebases have been curated, including OncoKB(1), My Cancer Genome(3), CIViC (4), Precision Medicine Knowledge Base(PMKB) (5), the JAX-Clinical Knowledgebase (CKB) (6), and Cancer Genome Interpreter (CGI) (7). Although clinically relevant, the interpretation of somatic variants is still not a standardized practice, and different clinical groups often generate different or even conflicting results. To standardize clinical interpretation of somatic variants in cancer and support clinical decision making, the Association for Molecular Pathology (AMP), American Society of Clinical Oncology (ASCO), College of American Pathologists (CAP), jointly proposed standards and guidelines for interpretation and reporting of somatic variants, which classify somatic variants into four Tiers: strong clinical significance (Tier I), potential clinical significance (Tier II), uncertain significance (Tier III), and benign (Tier IV) (8). The AMP/ASCO/CAP 2017 guideline included 12 pieces of evidence, which are diagnostic, prognostic and therapeutic clinical evidences, mutation types, variant allele fraction (mosaic variant frequency (likely somatic), non-mosaic variant frequency (potential germline)), population databases, germline databases, somatic databases, predictive results of different computational algorithms, pathway involvement, and publications (8,9).

However, since the AMP/ASCO/CAP classification scheme heavily relies on published clinical evidence for a variant, ambiguous assignments were still frequently observed among human curators, using the same evidence for a given variant. For example, Sirohi et al., compared human classifications for fifty-one variants by randomly selected 20 molecular pathologists from 10 institutions (10). The original overall observed agreement was only 58%. When providing the same evidential data of variants to the pathologists, the agreement rate of re-classification increased to 70%. The reasons for discordance are: (i) gathering information/evidence is quite complicated and may not be reproducible by the same interpreter at different time points; (ii) different researchers may prefer to use different algorithms, cutoffs and parameters, making the interpretation less reproducible; (iii) newly published evidence for certain variants might not been incorporated into the evaluation system instantly and systematically, which is especially relevant to the variants with unknown significance (VUS).

To standardize the interpretation of somatic variants across multiple knowledgebases, a more recently published knowledgebase, MetaKB from The Variant Interpretation for Cancer Consortium (VICC), aggregated evidences based on AMP/ASCO/CAP 2017 guideline (11). However, this meta knowledgebase also has the following limitations: 1) it only focused on consensus interpretations on a limited number of known ‘hotspot’ mutations, so that a large number of variants were currently classified as unknown clinical significance but may still play oncogenic roles through “loss of function” or “activating function” in cancer; 2) it only provided summarized classification for each variant, without demonstrating itemized evidence in details for each individual variant when mapping to the 12 criteria of AMP/ASCO/CAP 2017 guidelines; therefore, users cannot conduct customized evaluations based on their own protocols and experiences; and 3) it utilized a simple score system to rank driver mutations without considering heterogeneity of functional consequence (deleteriousness) of the variants, especially for those newly identified variants reported in publication.

In clinical practice, when a somatic mutation is considered to have strong confidence in causing functional impact on protein changes, clinicians likely interpret it as clinically significance or likely clinical significance (12,13). Although a number of remarkable software tools such as SIFT(14), Polyphen2 (15) and FATHMM(16) were developed to predict functional impacts, disagreements on certain mutations were consistently observed across these tools. Although later some meta-analysis tools such as DANN (17) and DriverPower (18) were developed to prioritize functionally important variants using more comprehensive functional scoring features as the input, they face the limitation in jointly modeling clinical impact features based on the AMP/ASCO/CAP guidelines. Because the guidelines tend to be conservative (“negative diagnosis” is preferred over “wrong diagnosis”), resulting in more than expected variants were misinterpreted as VUS (19-27). In addition, the AMP/ASCO/CAP guidelines only designated 7 functional impact prediction tools such as MutationAssessor (28) as the official recommended tools, and only the variant from majority voting (more than 4 from 7 tools) can be considered clinical significance, it over-simplified the heterogeneous functional consequence of variant in cancer development. While it may be useful in prediction of overall impact of driver genes, it is not optimal to prioritize novel variants found in the genes. To address these challenges and improve automated clinical interpretations for cancer variants, there is a strong need in the development of reliable and accurate computational methods, by utilizing both clinical evidence and functional impact score features.

We have previously developed a standalone software VIC written in Java, which was among the first tools to interpret clinical impacts of somatic variants using a rule-based scoring system based on 12 criteria of the AMP/ASCO/CAP 2017 guideline (29). In the current study, we developed an improved somatic variant interpretation tool called CancerVar implemented in Python (https://github.com/WGLab/CancerVar), with an accompanying web server (https://cancervar.wglab.org/). Compared to VIC, CancerVar is an advanced tool providing more options to users: (1) Python implementation provides more flexibility to incorporate CancerVar into custom command line workflows, (2) a user-friendly web server with pre-computed clinical evidence for 13 million variants coming from 1,911 cancer census genes through literature mining and database aggregations, (3) flexible AMP/ASCO/CAP rule-based score system and deep learning-based score system using semi-supervised generative adversarial network (SGAN) method to allow improved interpretations, (4) RESTful API to allow program developers to freely access complied knowledge. CancerVar allows users to query clinical interpretations for variants using chromosome position, cDNA change or protein change, and interactively fine-tune weights of scoring features based on their prior knowledge or additional user-specified criteria. Importantly, the CancerVar web server can generate automated texts with summarized descriptive interpretations, such as diagnostic, prognostic, targeted drug responses and clinical trial information for many hotspot mutations, which will significantly reduce the workload of human reviewers and advance the precision medicine in clinical oncology.

## MATERIAL AND METHODS

### Overview of clinical evidence mapping to the AMP/ASCO/CAP 2017 guidelines

According to the AMP/ASCO/CAP 2017 guidelines, there are a total of 12 types of clinical-based evidence to predict the clinical significance for somatic variants, including therapies, mutation types, variant allele fraction (mosaic variant frequency (likely somatic), non-mosaic variant frequency (potential germline)), population databases, germline databases, somatic databases, predictive results of different computational algorithms, pathway involvement, and publications (8,9). As shown in **Figure 1**, CancerVar contains all the above 12 evidence, among which 10 of them are automatically generated and the other two, including variant allele fraction and potential germline, require user input for manual adjustment.

**Figure 1.**
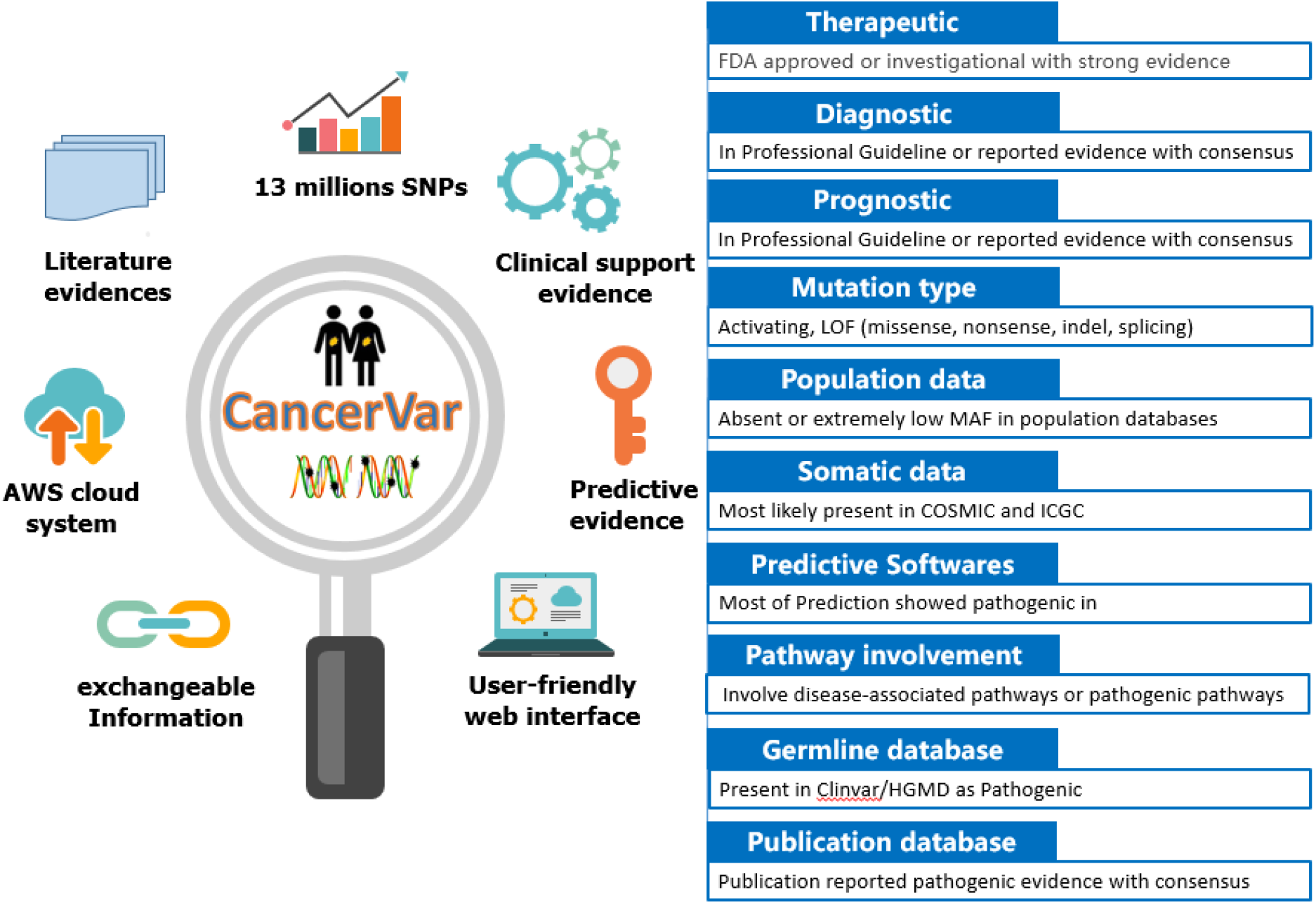
The functions of CancerVar and the descriptions of 12 types of evidence.

### Cancer variants collection and pre-processing

The cancer census gene list or potential driver gene list were very essential to all the somatic variants annotators. We curated a list of 1,911 cancer census or driver genes with 13 million exonic variants from 7 existing cancer knowledgebase, including COSMIC, CIViC, OncoKB, etc, and 2 datasets collected from literature about driver genes predictions (**Supplementary Table 1**). For each exon position in these 1,911 genes, we generated all three possible nucleotide changes. CancerVar fully scanned all the potential variants of significance, and it overcomes the limitations of other knowledgebase annotation datasets which only compiled variants reported or documented previously. We pre-compiled clinical evidence based on 2017 guideline for all the possible variant changes, which makes the variant searching in CancerVar very fast. In CancerVar, we documented all types of clinical evidence such as *in-silico* prediction, drug information, and publications in detail to help users making their own clinical decisions according to their prior knowledge.

### Evidence-based scoring method to prioritize clinical significance of somatic variants

CancerVar evaluates each set of evidence and scores each piece of clinical-based prediction (CBP). The variant evidence will get 2 points for strong clinical significance evidence or oncogenic,1 point for supporting clinical significance or oncogenic, 0 for no support, -1 for benign or neutral. The CancerVar score will be the sum of all the evidence. The complete score system for each CBP can be found in **Supplementary Table 2**. Let the *CBP*[*i*] be the *i* ^th^ evidence score, weight [*i*] is the score for *i* ^th^ evidence. The CancerVar score can be calculated in Equation 1. The weight is 1 by default, but users can adjust it based on its importance from prior knowledge. Based on the score range in Equation 2, we classify each variant into one of the four Tiers: strong clinical significance, potential clinical significance, unknown clinical significance (VUS), and benign/likely benign (neutral).

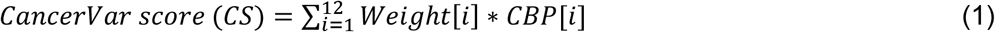

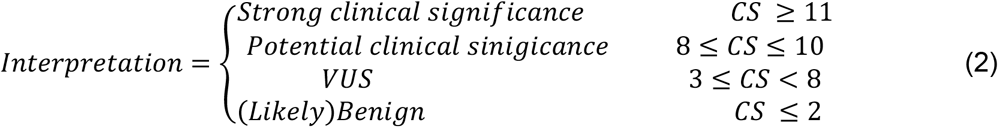

### Semi-supervised generative adversarial network (SGAN) to predict driver mutations

We developed semi-supervised generative adversarial network (SGAN) method to predict driver mutations using 12 clinical evidence prediction scores and 19 pre-computed scores predicted by other computational tools. The 19 predictive tools include: (1) nine function-prediction method: FATHMM (30), FitCons (31), MutationAssessor (32,33), MutationTaster (34), PolyPhen2-HDIV, PolyPhen2-HVAR (35), PROVEAN (36), SIFT (14), and VEST3 (37); (2) five ensemble methods: CADD (raw score and Phred score) (38), DANN (17), FATHMM-MKL (39), MetaLR (40), and MetaSVM (40); and (3) five conservation methods: GERP++ (41), PhastCons (42) (on vertebrate and mammalian separately), PhyloP (43) (on vertebrate and mammalian separately), LRT (44), and SiPhy (45). Since the score range are very diverse among the predictive tools, we used their categorical outputs as the prediction features. For some variants missed certain predictive values, we excluded the variants with more than 2 missing features. After filtering, a total of 12.9 million variants were used for downstream analysis. **Supplement Table 3** shows the distribution of missing rate before and after filtering and the maximum missing ratio is less than 10%.

Then, we applied SGAN artificial neural network to predict the probability of clinical significance after imputation. As shown in **Figure 2**, the SGANs architecture was originally developed in the context of unsupervised learning, which consists of 2 parts: generator and discriminator. The Generator (G) is to generate synthetic samples (fake) by random noise from the normal distribution; and the Discriminator (D) is to differentiate realistic samples and synthetic data. The Generator contains 3 linear layers with batch normalization, LeakyReLu as activation layer, and 60% dropout rate in each layer. The final layer is a linear layer with batch normalization and Tanh as activation layer. As for the discriminator (D), we implemented 3 CNN layers with Tanh as activation layers. The semi-supervised GAN is particularly useful in prediction of a huge amount of unlabelled samples using a small number of labelled samples. During training process, the SGAN learn the underlying distribution (clusters) of data samples by discriminating the synthetic samples and unlabelled realistic samples in each epoch, and meanwhile the network labels categories for clusters by classifying the labelled realistic samples. In this process, the discriminator/ classifier will be trained to discriminate the fake/real samples and to classify the labelled samples.

**Figure 2.**
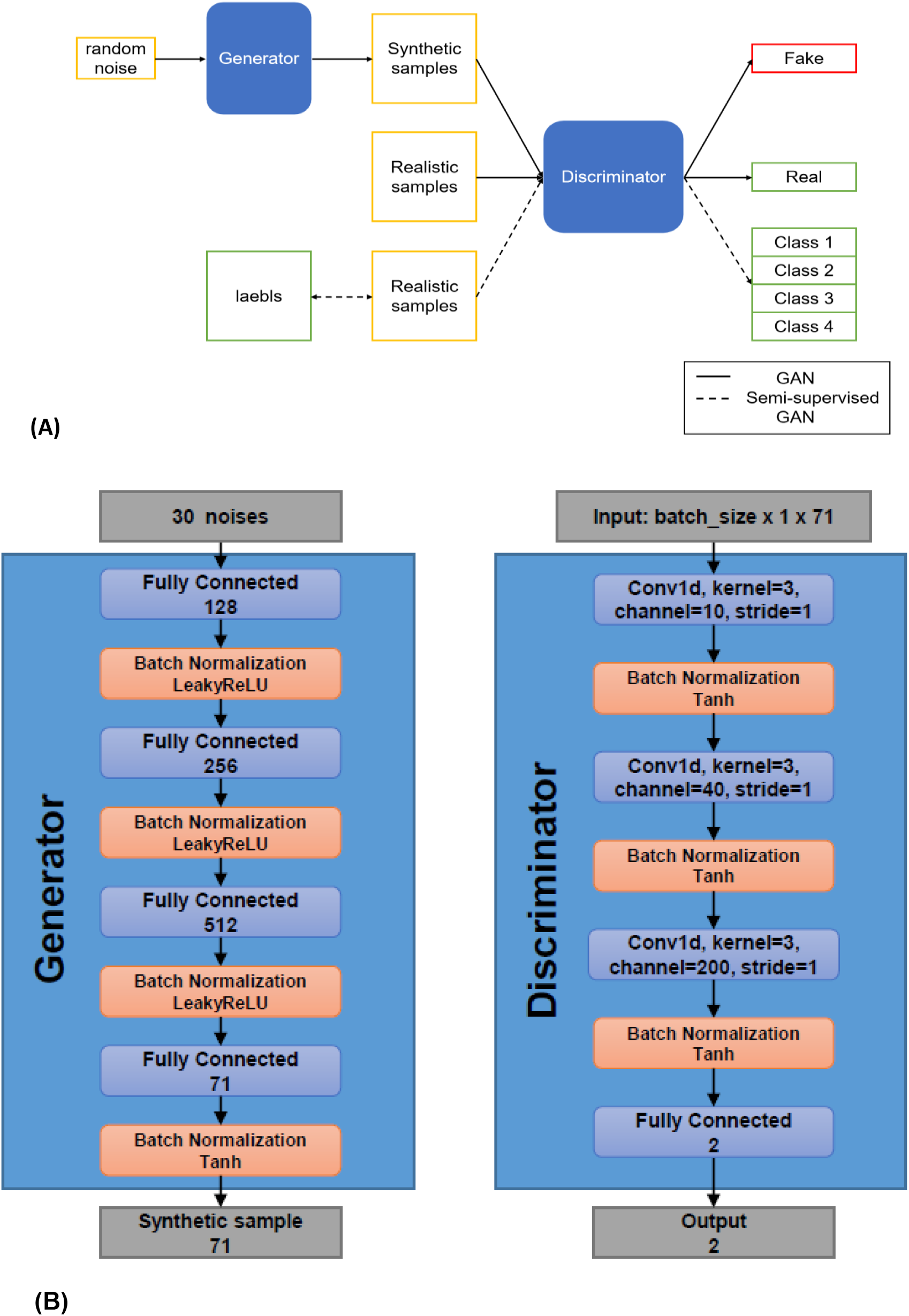
(A) Workflow of the SGAN method and (B) Architecture of generator and discriminator/classifier used in SGAN. Here we used a 3 transposed CNN layer to generate synthetic samples from a vector consisting of 100 random noises. The discriminator/classifier is a typical resNet-18.

In our semi-supervised GAN model, the input data consists of labelled samples, unlabelled samples and random noises from normal distribution. Firstly, the noises are converted to synthetic samples by the generator. Secondly, the discriminator/classifier classifies the sample into 3 classes: 1. neutral, 2. non-neutral (driver mutations), and 3. fake synthetic data, in which the unlabelled real samples can be identified as 1 or 2 and the synthetic samples is 3. Therefore, our models take in X as input and output a vector (*l*_1_, *l*_2_, *l*_3_) which can be converted to probability by Softmax function. As for supervised learning on labeled samples, the probability is: 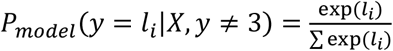 and we used *P*_*model*_ (*y* = 3|*X*) to infer the probability that X is fake. Meanwhile, the probability that X is real but unlabelled is 1 *− P*_*model*_ (*y* = 3|*X*). Therefore, the loss function L of our discriminator/classifier can be written as two parts:

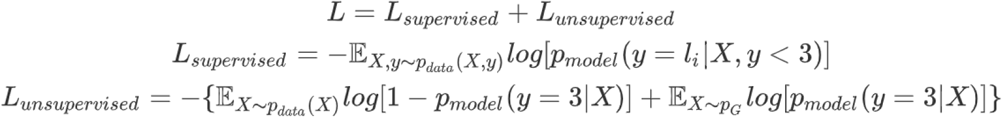

The *p*_*data*_ is the underlying distribution of real samples and *p*_*G*_ is the distribution of the output from generator. As for the loss of Generator, we used feature matching (46) as our loss function: 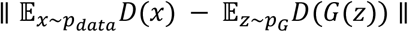.

### SGAN training and testing process

We implemented SGAN by PyTorch. For unlabelled data, we randomly selected 60,000 variants from 12.9 million samples with non-missing features. The evidence-based scores were converted into dummy features and added Gaussian noise (mean=0, std=0.02) to make the features continuous. The labelled data are from cancer patients of CHOP cancer cohort; we have 4,000 variants (1,000 are positive) as the training set and 1,234 variants (669 are positive) as the validation set. We tested the SGAN model on 6,226 variants (1,335 are positive) which were manually compiled from literature review. The missing feature values were filled with the mean value of the non-missing from its 40 nearest neighbouring variants in the whole training data by KNNImputater, a python package from scikit-learn (47).

As for the synthetic samples, the Generator generates random noise from standard normal distribution in each batch step, and outputs the synthetic samples. In each minibatch, the model calculates 2,000 labelled samples, 10,000 unlabelled samples and 10,000 synthetic samples from generator. The discriminator/classifier is trained by calculating the loss from supervised learning and unsupervised training separately. And then the generator is trained by minimizing the feature matching in each batch.

### Pan-Cancer Benchmarks from public dataset

Most of the somatic variant annotators or datasets have not been systematically assessed on their performance, especially for literature or knowledge-based tools. For systematic performance benchmarking of CancerVar, complementary and comprehensive benchmark datasets are needed and established for clinical significance prediction somatic variant. To robustly assess the performance of CancerVar, we employed several different benchmark datasets:(i) Multi-Institutional evaluation study (10) with fifty-one variants from Sirohi et; (ii)Literature annotation database from OncoKB (1) and CIViC(4); (iii) TP53 mutations on their target transcription activity from IARC database (48); (iv)Functional annotation based on in vitro cell viability assays from study of Ng. et al (49).

### Benchmarks from CHOP internal dataset

Importantly, other than datasets from public resource, we also have our internal dataset with 7967 somatic mutations from cancer patient’s cohort at the Children’s Hospital of Philadelphia (CHOP cancer cohort). Each variant has been manually annotated and classified by human experts in the diagnostic labs. Using the AMP/ASCO/CAP guideline, the 4-Tier classification assignment for each somatic mutation has to be agreed by at least two cancer experts from the Division of Genomic Diagnostics Lab at CHOP. Furthermore, to train the deep learning model, we used variants from strong clinical significance (Tier I) and potential clinical significance (Tier II) categories as the positive samples, and used variants from benign/likely benign (Tier IV) as the negative samples. In total, we have 5234 variants, in which 1668 are positive samples and 3566 are negative samples for training and validation in the SGAN model.

## RESULTS

### Summary functions of CancerVar

CancerVar provides multiple query options at variants-, gene-, and CNA levels across 30 cancer types and two versions of reference genomes: hg19 (GRCh37) and hg38 (GRCh38). Given user-supplied input, CancerVar generates an output web page, with information organized as cards including free text interpretation summary, gene overview, mutation information, evidence overview, pathways, clinical publications, protein domains, *in silico* predictions, exchangeable information from other knowledgebases. The CancerVar web server provides full details on the variants, including all the automatically generated criteria, most of the supporting evidence and predictive scores for clinical significance. CancerVar web service can be accessed at http://cancervar.wglab.org and the command-line program can be downloaded at https://github.com/WGLab/CancerVar.

Using rule-based approach, users have the ability to manually adjust these criteria and perform re-interpretation based on their prior knowledge or experience. If the user already know the information of each of the scoring criteria for the variant (possibly inferred by themselves using other software tools), they can alternatively compute the clinical significance of the variant from the “Interpret by Criteria” service instead. Each variant will be provided with a prediction score and clinical interpretation as strong clinical significance, potential clinical significance, uncertain significance, and likely benign/benign based on the 12 criteria of the AMP/ASCO/CAP guideline. Using deep learning-based approach, CancerVar provides probability score predicted by SGAN to determine the oncogenicity of a variant, using 12 evidence features from AMP/ASCO/CAP guidelines and computational metrics predicted by 19 tools.

### Performance assessment and comparative evaluations of CancerVar with external human manual annotator

Sirohi et. al measured the reliability of the 2017 AMP/ASCO/CAP guidelines (10) using fifty-one variants (31 SNVs, 14 indels, 5 CNAs, one fusion) based on literature review. Among these variants, we selected 43 variants including all 31 SNVs and 12 insertion-deletion variants (we did not find alternative alleles information for two indels in gene *CHEK1* and *MET*). CancerVar interpreted these 43 variants with the specified cancer types. Since these 43 variants do not have solid/consistent clinical interpretation, we compared 20 pathologists’ opinions from 10 institutions with CancerVar’s predictions. As shown in **Table 1**, CancerVar assigned 21 variants as Tire I/II (strong or potential clinical significance). Among these 21 variants, the pathologists classified 17 variants (17/21, around 81%) as Tire I/II in agreement. Moreover, CancerVar assigned 21 variants as VUS; among these 21 variants, 9 variants (9/21, around 43%) also be classified as VUS by pathologist reporters. In total, 26 variants (around 61%) have a match of clinical significance between human reporters and CancerVar.

**Table 1.** Comparison of classification on 43 variants between 20 pathologists and CancerVar.

| Annotators | Classifications | 20 pathologists |  |  |  |
| --- | --- | --- | --- | --- | --- |
|  |  | I/II* | III | IV | Total |
| CancerVar | I/II | 17 | 4 | 0 | 21 |
|  | III | 12 | 9 | 0 | 21 |
|  | IV | 0 | 1 | 0 | 1 |
|  | Total | 29 | 14 | 0 | 43 |

\* Tire I: strong clinical significance; Tire II: potential clinical significance; Tire III: unknown clinical significance (VUS); Tire IV: benign/likely benign

The interpretation details of these 43 variants can be found in the **Supplementary Table 3** and **Figure 3**. Compared to human interpreters, the advantage of CancerVar is clear in that it can automatically generate clinical interpretations with standardized, consistent and reproducible workflow, with evidence-based support for each of the 12 criteria. Therefore, CancerVar will greatly reduce the workload of human reviewers and facilitate the generation of precise and reproducible clinical interpretation.

**Figure 3.**
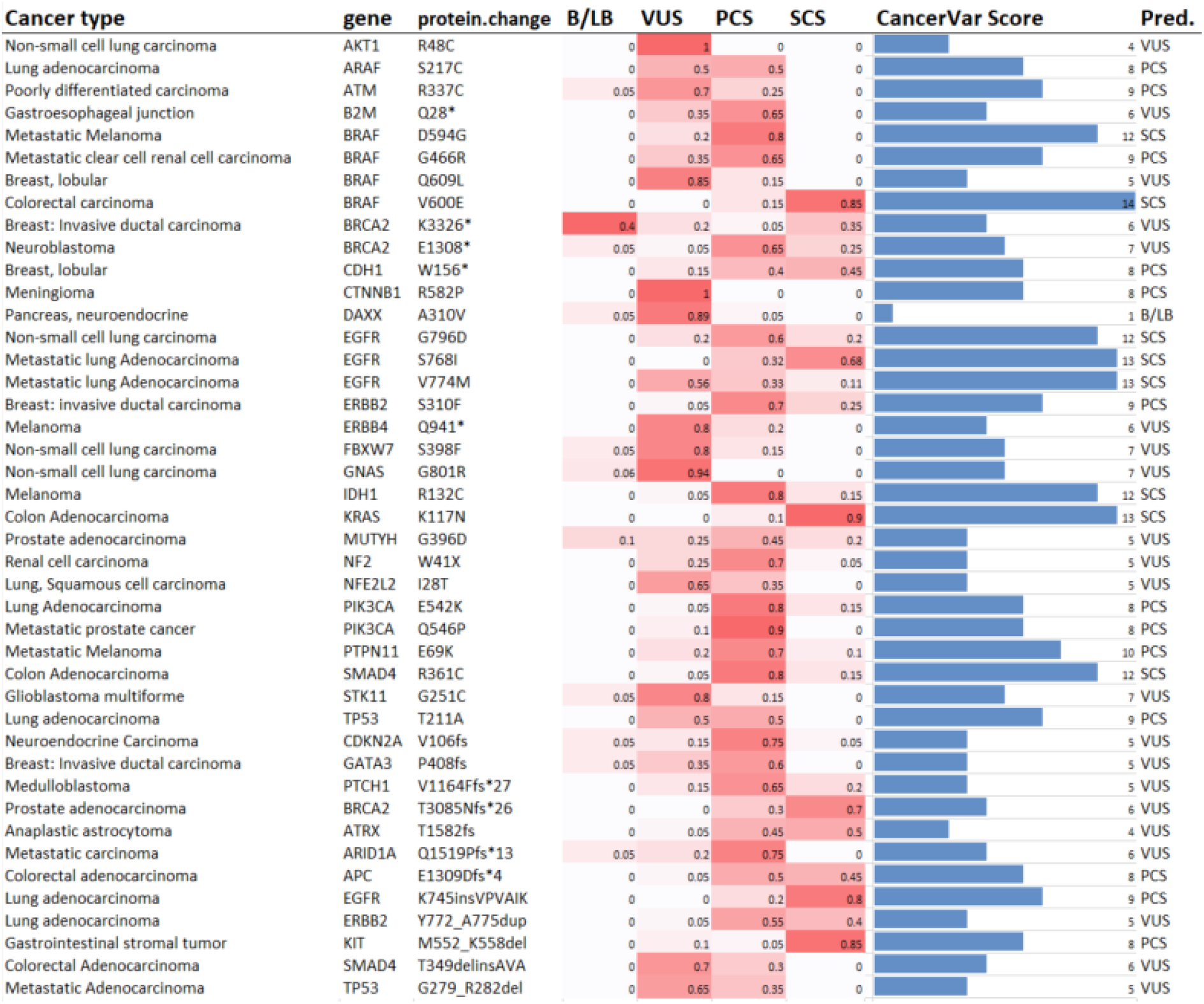
Comparison of interpretation on 43 variants between 20 pathologists and CancerVar. The heatmap shows the ratio of 20 pathologists voting for the four Tiers: Tier I strong clinical significance (SCS), Tier II potential clinical significance (PCS), Tier III variant uncertain clinical significance (VUS), and Tier IV benign/likely benign (B/LB). The last two columns are cancerVar predicted score and classification. Results show CancerVar has 81% (17/21) agreement rate with pathologists’ majority voting for Tier I/II, and 60.5% (26/43) agreement rate for all Tiers. This agreement rate is comparable to the 58% agreement rate within the 20 pathologists, but CancerVar can automate interpretation.

### OncoKB annotation benchmark

OncoKB (1), a manually curated database of cancer mutations oncogenic effect, has been widely used in cancer research community. OncoKB provides evidence classification system to interpret the genomic alterations and classified variants as inconclusive, likely neutral, Predicted oncogenic, likely oncogenic, or oncogenic. Totally, 3455 SNVs in 245 genes were downloaded from OncoKB annotation database (downloaded Mar/01/2020). This version contained 2582 oncogenic/likely oncogenic (O/LO) mutations, 587 likely neutral mutations, and 286 mutations annotated as inconclusive for this study. CancerVar evidence-based and DL-based prediction methods were applied to classify compiled mutations and compared with OncoKB classifications. For the O/LO group in the OncoKB, CancerVar rule-based method classified 1839 (1839/2582, 71.2% consistent with OncoKB classification) variants as strong or potential clinical significance; while CancerVar deep learning-based method classified 2319 variants (2319/2582, 90% consistent with OncoKB classification). The details are in **Table 2 and Figure 4(a)**. The UpSet plot showed the prediction intersections between OncoKB, CancerVar rule-based and deep learning-based methods.

**Figure 4.**
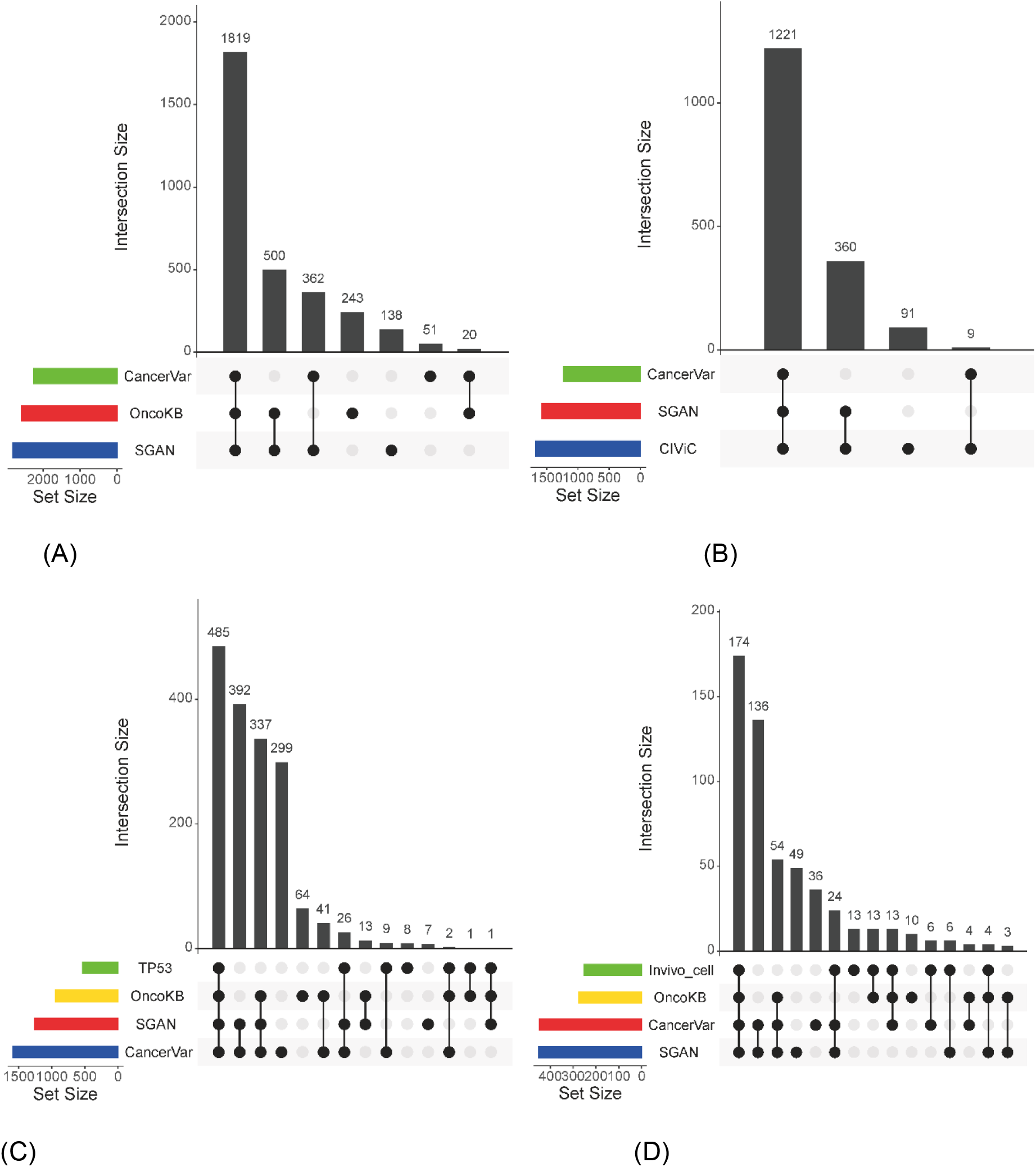
UpSetR (Conway et al., 2017) plot highlights the intersection of multiple methods with oncogenic prediction from different datasets. (A) Mutations were taken from OncoKB dataset. (B) Mutations were taken from CIViC. (C) Mutations were taken from IARC TP53 Transactivation dataset. (D) Mutations were taken from Cell Viability in Vitro by Ng. et al, 2018

**Table 2.**
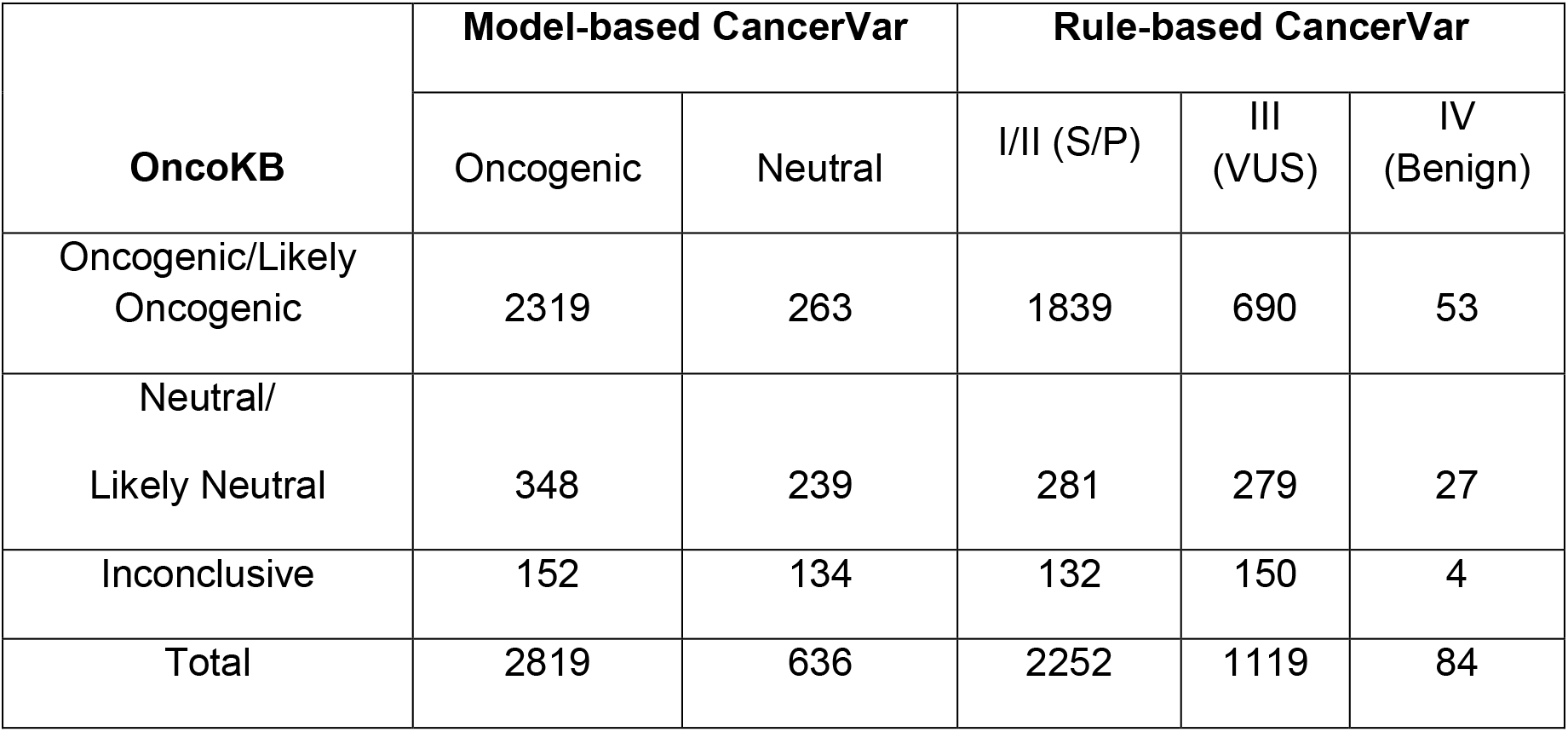
Summary of CancerVar prediction on OncoKB mutations.

### CIViC annotation benchmark

CIViC is a crowd-sourced and expert-moderated public resource for somatic variants in cancer (4). It adopts five evidence levels to differentiate reported mutations, namely A: validated, B: clinical, C: case study, D: preclinical, and E: inferential. In total, 1681 unique SNVs/INDELs from 113 unique genes were retrieved from CIViC website (https://civicdb.org/releases, accessed in May/01/2020) and assessed by the CancerVar program. CancerVar rule-based method predicted 1230 (1230/1681, 73.2% consistent with CIViC classification) variants as strong or potential clinical significance, while CancerVar DL model-based method predicted 1581 (94.1% consistent with CIViC classification). **Table 3 and Figure 4(b)** have the details of CancerVar prediction.

**Table 3.** Summary of CancerVar prediction on CIViC mutations.

| **CIViC** | **Model-based CancerVar** |  | **Rule-based CancerVar** |  |  |  |
| --- | --- | --- | --- | --- | --- | --- |
|  | Oncogenic | Neutral | I (Strong) | II (Potential) | III (VUS) | IV (Benign) |
| A: validated | 17 | 2 | 2 | 7 | 0 | 10 |
| B: clinical | 259 | 40 | 111 | 109 | 61 | 18 |
| C: case study | 792 | 30 | 178 | 439 | 198 | 7 |
| D: preclinical | 466 | 22 | 91 | 277 | 119 | 1 |
| E: inferential | 47 | 6 | 5 | 11 | 33 | 4 |
| Total | 1581 | 100 | 387 | 843 | 411 | 40 |

### IARC TP53 Transactivation mutation benchmark

TP53 is the most frequently mutated gene in human cancers, its mutants had been functionally assessed based on the median transactivation levels and complied as IARC TP53 database (48). Based on the median of 8 different yeast functional assays (WAF1, MDM2, BAX, h1433s, AIP1, GADD45, noxa, and P53R2), the TP53 mutations can be classified as lower transactivation (a median transactivation level <= 25% wild type) as oncogenic and higher transactivation (level >= 25% wild type) as neutral. We retrieved 1915 missense mutations (532 mutations were used as oncogenic cases and 1383 mutations were used as neutral cases) from this IARC TP53 database. For 532 oncogenic mutations in IARC TP53 database, CancerVar rule-based method predicted 522 (TP=98%) variants and model-based method predicted 512 (TP=96.2%) variants as strong/potential clinical significance. Compared to OncoKB predicted 489 (489/532=91.9%) variants, CancerVar rule-based method has a higher true positive rate. The details of CancerVar and OncoKB prediction can be viewed in **Table 4 and Figure 4(c)**.

**Table 4.** CancerVar and OncoKB predictions of mutations in the IARC TP53 transactivation dataset.

| **IARC TP53**<br>**Transactivation** | **Model-based**<br>**CancerVar** |  | **Rule-based CancerVar** |  |  | **OncoKB** |  |  |
| --- | --- | --- | --- | --- | --- | --- | --- | --- |
|  | Oncogenic | Neutral | S/P | VUS | benign | O/LO | VUS | Neutral |
| Oncogenic | 512 | 20 | 522 | 10 | 0 | 489 | 43 | 0 |
| Neutral | 749 | 634 | 1069 | 314 | 0 | 455 | 928 | 0 |
| Total | 1261 | 654 | 1591 | 324 | 0 | 944 | 971 | 0 |

### Cell Viability in Vitro Assay benchmark

The oncogenic effect of somatic mutation can be directly assessed by preferential growth or survival advantage to the cells using some cellular assays. Ng et al. recently developed a medium-throughput in vitro system to test functional effects of mutations using two growth factor dependent cell lines, Ba/F3(a sensitive leukaemia cell line, frequently used in drug screening) and MCF10A(a breast epithelial cell line)(49). The cell viability data of mutations in these two cell lines were used to generate consensus functional annotation to distinguish mutations. The mutations were considered as oncogenic when cell viability was labelled as activating and as neutral when cell viability was labelled as neutral from the consensus functional annotation. Finally, we retrieved 717 missense mutations (253 as oncogenic, 464 as neutral) in 44 genes. In 253 oncogenic variants, CancerVar rule-based method predicted 217 (TP=85.7%) variants and DL model-based predicted 208 (82.2%) as strong/potential clinical significance, while OncoKB predicted 204 (TP=80.6%) variants as oncogenic, likely or predicted oncogenic (**Table 5 and Figure 4(d)**). Still, CancerVar rule-based method performs better than OncoKB.

**Table 5.** CancerVar and OncoKB predictions of mutations in the *in vitro* Cell Viability dataset by Ng. et al, 2018.

| **Cell Viability**<br>**(*in vitro*)** | **Model-based**<br>**CancerVar** |  | **Rule-based**<br>**CancerVar** |  |  | **OncoKB** |  |  |
| --- | --- | --- | --- | --- | --- | --- | --- | --- |
|  | Oncogenic | Neutral | S/P | VUS | benign | O/LO | VUS | Neutral |
| Oncogenic | 208 | 45 | 217 | 34 | 2 | 204 | 39 | 10 |
| Neutral | 242 | 222 | 230 | 222 | 12 | 71 | 335 | 58 |
| Total | 450 | 267 | 447 | 256 | 14 | 275 | 374 | 68 |

### SGAN performance for oncogenic variant’s prediction

We used cuda to accelerate the training process, which took ∼100 hours to train the model to 1000 epochs with a Nvidia Tesla M40 GPU card. SGAN model can learn the hidden distribution of unlabeled mutations comparing with the prediction in clinical data from the model trained only with labeled data. SGAN was compared with other six machine learning algorithms including gradient boosting tree (GBDT), support vector machine (SVM), AdaBoost (ADA), multi-layer perceptron (MLP), random forest (RF) and majority voting classifier (VC), which were discussed in a recently published paper AI-driver for driver mutation prediction in cancer (50). We further compared the performance with the other nine score schemes including LRT (44), DANN (17), CADD (38), FATHMM (30), SIFT (14), and MetaSVM (40), using area under the curve (AUC) score from receiver operating characteristic (ROC) plots and true negative rate (TNR, or specificity) as measurements. According to the performance evaluation on the independent testing set of 6226 somatic variants, **Figure 5** shows that CancerVar SGAN method (AUC=0.8595) performs the best compared to cancer-specific driver predicting methods and any individual functional prediction tool.

**Figure 5.**
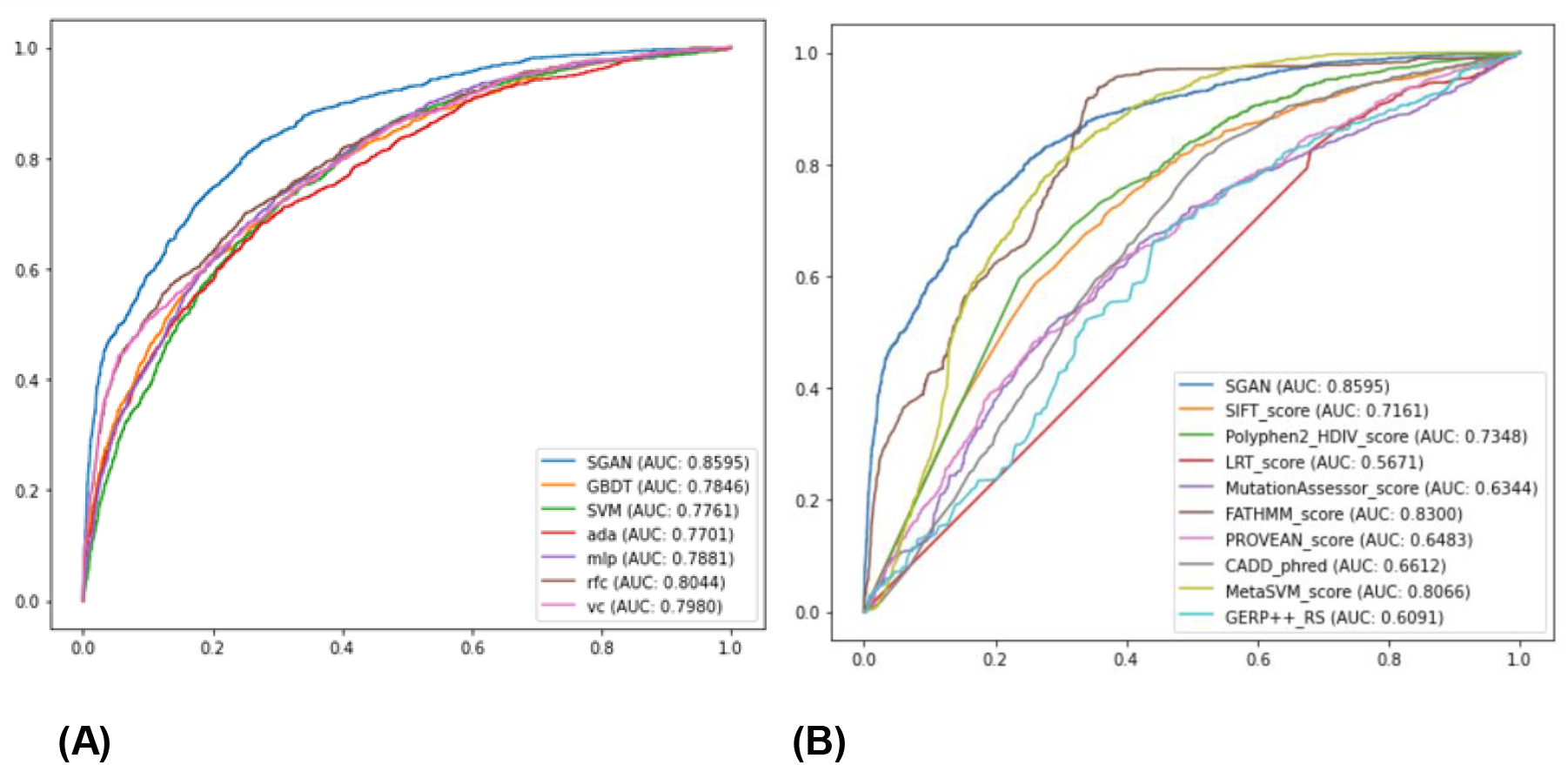
Receiver operating characteristic (ROC) curves for performance comparison on 6,226 somatic mutations as the testing set. (A) SGAN outperforms six other machine learning algorithms including gradient boosting tree (GBDT), support vector machine (SVM), AdaBoost (ADA), multi-layer perceptron (MLP) random forest classification (RFC), and voting classifier (VC). (B) SGAN also outperforms any individual functional impact prediction tool in prediction of cancer somatic driver mutations.

### FDA approved or recognized cancer biomarkers for therapeutic, diagnosis and prognostic

To show the performance and reliability, we also collected 22 cancer biomarkers approved by the US Food and Drug Administration (FDA, then interpreted these biomarkers and predicted their oncogenicity using our SGAN model. In these 22 biomarkers, 9 of them were classified as Tier-I strong clinical significance and rest of 13 were classified as Tier-II potential clinical significance when using only evidence. For the SGAN model based on deep learning, most of the biomarkers (19 out of 22) were predicted with score >=0.95 means higher probability as oncogenic. The interpretation of these biomarkers are showed in the **Table 6**.

**Table 6.** FDA approved or recognized biomarkers (therapeutic, diagnosis and prognostic) clinical significance or oncogenic prediction from CancerVar and SGAN.

| Gene | Alternation | Cancers | Levels | CancerVar | SGAN |
| --- | --- | --- | --- | --- | --- |
| ABL1 | T315I | B-Lymphoblastic Leukemia/Lymphoma/Chronic Myelogenous Leukemia | Therapeutic (Ponatinib) | Tier-II(score 8) | 0.87 |
| AKT1 | E17K | Breast Cancer/Ovarian Cancer/Endometrial Cancer | Therapeutic (AZD5363) | Tier-II(score 10) | 0.97 |
| BRAF | V600E | Melanoma/Non-Small Cell Lung Cancer | Therapeutic (Dabrafenib + Trametinib; Vemurafenib) | Tier-I(score 11) | 0.98 |
| EGFR | T790M | Non-Small Cell Lung Cancer | Therapeutic (Osimertinib) | Tier-II(score 9) | 0.96 |
| EGFR | L861Q | Non-Small Cell Lung Cancer | Therapeutic (Afatinib) | Tier-II(score 10) | 0.98 |
| EGFR | S768I | Non-Small Cell Lung Cancer | Therapeutic (Afatinib) | Tier-I(score 11) | 0.99 |
| EZH2 | Y646F | Follicular Lymphoma | Therapeutic (Tazemetostat) | Tier-I(score 11) | 0.99 |
|  | Y646H |  |  | Tier-I(score 11) | 0.99 |
|  | Y646N |  |  | Tier-I(score 11) | 0.99 |
|  | Y646S |  |  | Tier-I(score 11) | 0.99 |
| FGFR3 | R248C | Bladder Cancer | Therapeutic (Erdafitinib) | Tier-II(score 9) | 0.99 |
|  | S249C |  |  | Tier-II(score 9) | 0.99 |
|  | Y373C |  |  | Tier-II(score 10) | 0.99 |
| JAK2 | V617F | Primary Myelofibrosis | Prognostic | Tier-I(score 11) | 0.83 |
| KIT | A829P | Gastrointestinal Stromal Tumor | Therapeutic (Imatinib; Regorafenib; Ripretinib; Sunitinib) | Tier-II(score 9) | 0.86 |
|  | T670I |  |  | Tier-II(score 9) | 0.97 |
|  | V654A |  |  | Tier-I(score 11) | 0.99 |
|  | Y823D |  |  | Tier-II(score 10) | 0.99 |
|  | D816V | Systemic Mastocytosis | Diagnostic | Tier-II(score 9) | 0.98 |
| KRAS | G12C | Non-Small Cell Lung Cancer | Therapeutic (AMG-510) | Tier-I(score 11) | 0.99 |
| PDGFR A | D842V | Gastrointestinal Stromal Tumor | Therapeutic (Avapritinib) | Tier-II(score 9) | 0.95 |
|  | D842Y |  |  | Tier-II(score 10) | 0.99 |

### Use case: example of comprehensive interpretation of *FOXA1* somatic mutation in Prostate Cancer

In this use case, we showed the clinical interpretation of two mutations in prostate cancer (Figure 6) from rule-based and deep learning model. Prostate cancer is the most commonly diagnosed cancer in men in the world (51). The FOXA1 protein (Forkhead box A1, previously known as HNF3a) is essential for the normal development of the prostate (52). The *FOXA1* somatic mutations have been observed frequently in prostate cancer(53) and are associated with poor outcome. However, the mechanism of driving prostate cancer by mutations in *FOXA1* was still not clear. In 2019, two papers published in Nature demonstrated that *FOXA1* acts as an oncogene in prostate cancer (54,55). They found that the hotspot mutation at R219 (R219S and R219C) drove a pro-luminal phenotype in prostate cancer and exclusive with other fusions or mutations (54,55). We interpreted these two mutations, but here we only illustrated the clinical interpretation for R219C since the interpretation result of R219S was very similar to R219C. We searched this missense mutation R219S using protein change and gene name as “FOXA1” in the CancerVar web server. CancerVar did not find any therapeutic, diagnostic and prognostic evidence for this mutation. Since this mutation has been recently incorporated in somatic databases including COSMIC (ID: COSM3738526) and ICGC (ID: MU67448716), CBP_9 as moderate evidence applied. Recently two publications reported its biological functions in prostate cancer, CBP_12 applied. In addition, from CBP_7, this mutation is absent or has extremely low minor allele frequency in the public allele frequency database. All seven *in silico* methods predicted this mutation as (likely) pathogenic, CBP_10 applied. According the AMP/ASCO/CAP/CGC guidelines, this variant falls into the class of “Tier III uncertain significance” with a score of 7, but very closed the class of “Tire II potential”. While from deep learning of SGAN, the score is 0.99 as Oncogenic. This semi-automated interpretation approach can greatly improve the prediction accuracy for each variant, given existing knowledge and domain expertise, while, a model-based approach involving machine intelligence such as SGAN model still can be as another optimized option and could be explored in our future work.

**Figure 6.**
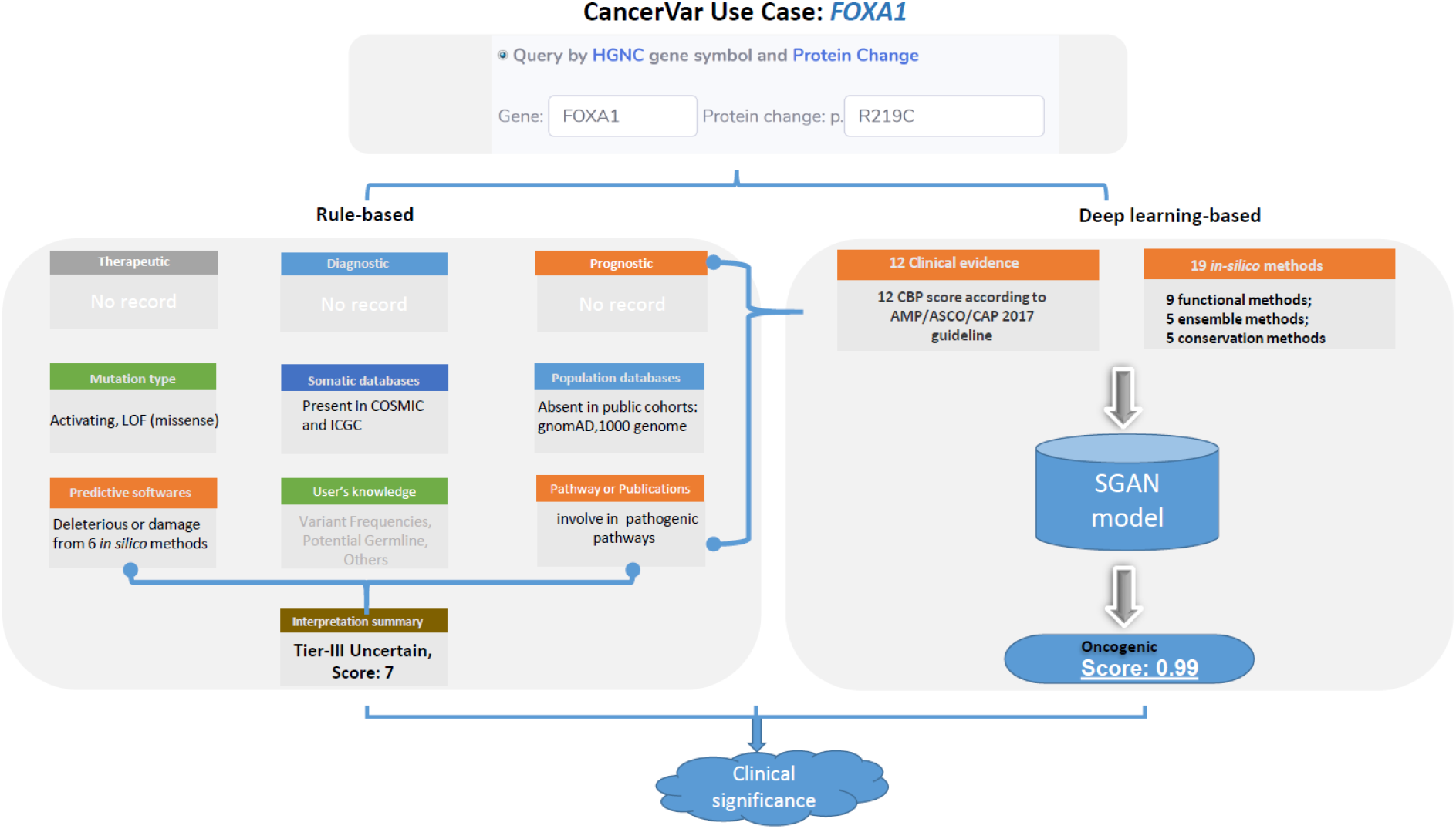
CancerVar use case of *FOXA1*. We queried FOXA1 mutation R219C in prostate cancer. The rule-based prediction of this variant was Tier-III uncertain-significance with score 7, very close Tier-II. However, after applied deep learning, the SGAN model predicted this variant as oncogenic with score 0.99. Finally, we suggested this variant with clinical significance.

## DISCUSSION

Clinical interpretation of cancer somatic variants remains an urgent need for clinicians and researchers working in the field of precision oncology, especially given the transition from panel sequencing to whole exome/genome sequencing in cancer genomics. To build a standardized, rapid and user-friendly interpretation tool, we developed a web server (together with command-line software tools) to assess the clinical impacts of somatic variants using the AMP/ASCO/CAP guidelines. CancerVar is an enhanced version of cancer variants knowledgebase incorporated from our previously developed tools for variant annotations and prioritizations including InterVar (56), VIC (9), iCAGES (57), as well as assembling existing variants annotation databases such as CIViC(4), CKB(6) and OncoKB(1). We stress here that CancerVar will not replace human acumen in clinical interpretation, but rather to generate evidence to facilitate/enhance human reviewers by providing a standardized, reproducible, and precise output for interpreting somatic variants.

In CancerVar, we did not reconcile the well-known “conflicting interpretation” issues across knowledgebases; instead, we documented and harmonized all types of clinical evidence (i.e. drug information, publications, etc) for both hotspot and non-hotspot mutations in detail to allow users make their own clinical decisions based on their own domain knowledge and expertise. Compared to existing knowledgebases such as OncoKB, CIViC and metaKB, CancerVar provides an improved platform in four areas: (i) comprehensive, evidence-based annotations with rigorous quality control for ∼13 million somatic variants, which is not limited to the small number of known hotspot mutations; (ii) well-designed, flexible scoring system allowing users to fine-tune the importance of clinical evidence criteria according to their own prior knowledge; (iii) improved prioritization for cancer driver mutations using novel semi-supervised DL learning method; (iv) automatically summarized interpretation text so that users do no need to query evidence from multiple knowledgebase manually. We expect CancerVar to become a useful web service for the interpretation of somatic variants in clinical cancer research.

We also need to acknowledge several limitations in CancerVar. First, the scoring weight system is not very robust. We note that the existing clinical guidelines did not provide the recommendations for weighting different evidence types, and therefore treated all weights as equal by default; however, with the increasing amounts of clinical knowledge on somatic mutations, we expect that we may build a weighted model in the future to enhance the prediction accuracy. Second, a small number of CNAs (similar to hotspot mutations) has emerged as important biomarkers for disease characterization and therapeutic decision making, however there is a lack of specific database for clinically actionable somatic CNAs. Although AMP/ASCO/CAP published a CNAs guideline recently, CNAs are very heterogeneous in size, so their significance is much harder to score in practice. Therefore, in the future, we will design and implement the scoring system for CNAs with AMP/ASCO/CAP team, based on the platform used to discover CNAs, the reliability of the CNA calls, the genes covered by the CNAs and additional cancer type specific information from existing databases (given that different cancer types have different CNA profiles). Third, CancerVar currently cannot interpret inversions and gene fusions, and cannot interpret gene expression alterations, even though these genomic alterations may also play important roles in cancer development/progression. Before a specific guideline for these types of mutations becomes available, we suggest that users treat them as CNAs (gene inversions/fusions as deletions, and gene expression down-regulation or up-regulation as deletions or duplications).

Accurate clinical significance interpretation depends greatly on the harmonization of evidence, which should be precisely derived and standardized from multiple databases and annotations. Compared to existing knowledgebases that document limited number of hotspot mutations, CancerVar provides polished, comprehensive, and semi-automated clinical interpretations for large scale somatic variants with completed clinical evidences, and it greatly facilitates human reviewers draft clinical reports for panel sequencing, exome sequencing or whole genome sequencing on cancer. Although some commercial software tools also used AMP/ASCO/CAP rule to standardize variants interpretation, they requires a high license fee that pushes many academic researchers away. Importantly, besides the interpretation based on AMP/ASCO/CAP human experts’ consensus rules, the CancerVar deep-learning based SGAN approach jointly modeled both rule-based clinical features and functional prediction features to support oncogenic predictions for mutations. We believe that CancerVar allows comprehensive clinical interpretations and prioritizations for both hotspot and non-hotspot variants, achieving a significant impact to facilitate the implementation of precision oncology.

In summary, CancerVar is both a web server and a command-line software that provide polished and semi-automated clinical interpretations for somatic variants in cancer. In addition, it facilitates drafting clinical reports semi-automatically for panel sequencing, exome sequencing or genome sequencing on cancer. We expect to continuously improve CancerVar and incorporate new functionalities in the future, similar to what we have done on the wInterVar server and wANNOVAR server.

## CancerVar software accessibility

Users can access CancerVar through three ways, including a web server that is free and open to all users without login requirements (http://cancervar.wglab.org), a command-line software written in Python that is freely available from GitHub (https://github.com/wglab/CancerVar) for non-commercial users, and a RESTful API service to facilitate other web developers to access our pre-computed evidence for 13 million variants.

## ACKNOWLEDGEMENT

We would like to thank Drs. Sebastiao N. Martins-Filho and Nhu-An Pham for constructive comments. We also thank members of the Wang lab for helpful comments on the user interface of the CancerVar web server and for testing the CancerVar web server.

## FUNDING

This work was supported by the National Institutes of Health (NIH)/National Library of Medicine (NLM)/National Human Genome Research Institute (NHGRI) [grant number LM012895] and National Institutes of Health (NIH)/National Institute of General Medical Sciences (NIGMS) [grant number GM120609 and GM132713], and CHOP Research Institute.

## CONFLICT OF INTEREST

KW indirectly own shares but is not involved in the operation of PierianDx, which develops cloud-based solution for clinical interpretation of somatic mutations.

## Notes

http://cancervar.wglab.org

### Competing Interest Statement

The authors have declared no competing interest.

### Summary of Updates

We made some changes in main text and added Artificial Intelligence parts in our model.

